## Supplementary Information for "Molecular resolution imaging by post-labeling expansion single-molecule localization microscopy (Ex-SMLM)"

**Supplementary Table 1.** Formulas and values used to simulate the molecular expansion factor of microtubules

| **Expansion method** **(experimentally determined peak-to-peak distance)** | **Inner radius** $r_1$ | **Outer radius** $r_2$ | **Expansion factor** $f_{exp}$ |
|---|---|---|---|
| ProExM (pre-labeled)* (137.1 nm) | $\frac{(25+2*8.75)f_{exp}}{2}$ | $\frac{(25+4*8.75)f_{exp}}{2}$ | 3.1 |
| ProExM (pre- and post labeled)* (79.5 nm) | $r_1pre = \frac{25f_{exp}}{2}$ $r_1post = \frac{(25+2*8.75)f_{exp}}{2}$ | $r_2pre = \frac{25f_{exp}}{2} + 2*8.75$ $r_2post = \frac{(25+4*8.75)f_{exp}}{2}$ | 3.2** |
| ExM protocol (using trifunctional Al532 DNA) (pre-labeled)* (226.7 nm) | $\frac{(25+4*8.75+2*6.5)f_{exp}}{2}$ | $\frac{(25+4*8.75+2*7.14)f_{exp}+2*7.14}{2}$ | 3.2 |
| ExM protocol using Cy5 DNA label (pre-labeled)* (226.7 nm) | $\frac{(25+4*8.75+2*1.84)f_{exp}}{2}$ | $\frac{(25+4*8.75+2*7.14)f_{exp}}{2}$ | 3.2 |
| ExM-GA protocol (pre-labeled)* (133.8 nm) | $\frac{(25+2*8.75)f_{exp}}{2}$ | $\frac{(25+4*8.75)f_{exp}}{2}$ | 3.0 |

*Pre- and post-labeling refers to the time point of immunostaining before or after expansion. The size of the fluorophores has not been taken into account.

**Expansion factor results from the assumption that the pre-expansion label is diluted 1:10 compared to the post-expansion label.

**Supplementary Note 1.** Theoretical basis of the assumed values for the simulation of peak-to-peak distances.

We used a cylindrical function y to simulate the theoretical intensity profile of microtubules for different expansion protocols:

$$y = \begin{cases} h(\sqrt{r_2^2 - (x-c)^2} - \sqrt{r_1^2 - (x-c)^2}), \text{ if } \|x\| < r_1 \\ h(\sqrt{r_2^2 - (x-c)^2}), \text{ if } \|x\| \geq r_1, \|x\| < r_2 \\ 0, \text{else} \end{cases}$$

with $r_1$ and $r_2$ denoting the inner and outer cylinder radius, $h$ the profiles intensity and $c$ the center of the microtubule.

*Unexpanded microtubules.* For unexpanded microtubules, we assumed a diameter of 25 nm for the structure itself and 60 nm for indirect immunolabeled microtubules decorated with primary and secondary antibodies[1]. The broadening caused by immunolabeling with primary and secondary antibodies is thus 17.5 nm and accordingly for a single antibody 8.75 nm. The inner and outer radius $r_1$ and $r_2$ used in the cylinder function is defined by the ring-like area in which the secondary antibody is located around the microtubule. The diameter of microtubules of 60 nm and the 8.75 nm value of antibody broadening were used as basis for the calculation of expanded radii.

*ProExM and ExM-GA protocol (pre-labeling ExM).* In proExM and ExM-GA expanded samples microtubules are immunolabeled pre-expansion with primary and fluorophore conjugated secondary antibodies. The values for $r_1$ and $r_2$ can be transferred from the formula used for unexpanded microtubules labeled with primary and secondary antibodies but multiplied by the respective expansion factor.

*ProExM (pre- and post-labeling).* For the simulation of intensity profiles of pre- and post-labeled microtubules, we used the sum of two cylindrical functions $c_{pre}$ and $c_{post}$ with radii $r_{1,pre}$ ,$r_{2,pre}$ and $r_{1,post}$ ,$r_{2,post}$, respectively. $c_{post}$ describes the intensity profile of the post-labelling signal with an inner radius $r_{1,post}$ defined by the expansion of the microtubule radius and the broadening of the primary antibody introduced post-expansion (12.5 nm * expansion factor + 8.75 nm). Here, we assumed that the secondary antibody attaches to the primary antibody and is able to orient to the center or to the outside of the microtubule. This assumption reduces the inner radius $r_{1,post}$ to 8.75 nm. The outer radius $r_{2,post}$ is determined by the broadening of the secondary antibody similar to pre-expansion broadening by antibodies with 8.75 nm (12.5 nm * expansion factor + 2 * 8.75 nm). $c_{pre}$ describes the intensity profile of the pre-labeling signal that is calculated with similar values as used in '*ProExM and ExM-GA protocol (pre-labeling)*' protocol. The signal of the pre-expansion label is included in the calculation considering the dilution factor of 1:10 ($3.2^2$) for the 2D projected signals compared to the post-expansion label (expansion factor$^2$).

*Expansion protocols using DNA-labels.* ExM protocols using DNA-labels use primary and fluorescently-labeled modified secondary antibodies. The inner and outer radius $r_1$ and $r_2$ is determined by the diameter of the microtubule, the broadening effect of primary and secondary antibody and the broadening caused by the length of the DNA molecule and the position of fluorophores within the oligo. (Supplementary Figure 7a). Since protocols using DNA-labels belong to pre-labeling ExM protocols the inner and outer radius is effected by the antibodies as described in '*ProExM and ExM-GA protocol (pre-labeling)'* with the further enlargement of the distance of fluorophores from the secondary antibody. Since DNA polymers arranged in a double helix extend 0.34 nm per base pair (bp), we calculated the distance of the fluorophores depending on their respective position in the dsDNA. Thus, for '*ExM protocol using Cy5 DNA label (pre-labeled)*' fluorophores are positioned evenly distributed along the DNA at a distance of 2.38 nm (7 base pairs) (Supplementary Table 1). For *'ExM protocol using trifunctional DNA (pre-labeled)*' fluorophores are set at base position 21 and 42 (5'→3') positioning the fluorophores 7.14 nm and 14.28 nm away from the antibody, respectively.

*ExM protocol using trifunctional Alexa Fluor 532 DNA (pre-labeled).* We first calculated the outer radius $r_2$ assuming a double-stranded DNA with a length of 14.28 nm arranged as fully stretched dsDNA and oriented to the outside of the microtubule. The position of the outer fluorophore multiplied with the respective expansion factor gives the outer radius $r_2$. Thus, the inner fluorophore located closer to the antibody influences the distance for the inner radius $r_1$.

Since the maximum angle in which the DNA can be attached to the antibody is unknown, we used experimentally determined data to describe the inner radius of the cylindrical function. We therefore adjusted the value of the inner radius to fit the peak-to-peak distance of 226.7 nm (**Fig. 2m**) which allowed us to calculate a feasible angle φ in which the DNA is oriented:

$$\varphi = \cos^{-1}\left(\frac{3.9\ \text{nm} * f_{exp}}{14.28\ \text{nm} * f_{exp} - 7.14\ \text{nm}}\right),$$

with 3.9 nm as additional broadening of the inner radius introduced by the DNA, deriving from experimental data,14.28 nm as length of the 42 bp DNA, and 7.14 nm as distance of the innermost fluorophore from the linkage position.

Thus, for an expansion factor of 3.2x we determined a possible angle between -57.4° and + 57.4° of the 42 bp long DNA strand, relative to the radial axis (Supplementary Figure 7b).

*ExM protocol using Cy5 DNA label (pre- labeled).* We used the minimal DNA strand angle of ± 57.4° to determine the expansion factor of samples expanded according the ‘DNA Cy5 protocol‘ with a peak-to-peak distance of 201.0 nm (Fig. 2q). Assuming a fully stretched dsDNA that is linked pre-expansion into the hydrogel would result in an expansion factor of 2.86x.

As ssDNA is a kind of flexible polymer, the persistence length is shorter. Based on our experimental data we determined a coiling factor of 50 % of the oligonucleotide that matches an expansion factor of 3.2x if the DNA-strand is arranged between ±57.4°:

$$r_1(unexpanded) += \cos(\varphi)\frac{7.14\text{ nm } f_{exp} - 11.9\text{ nm}}{f_{exp}},$$

with 11.9 nm as distance of the innermost fluorophore from the acrydite linker group in an uncoiled state after applying the oligonucleotide and 7.14 nm as linking position of the coiled DNA-strand.

**Supplementary Table 2.** DNA sequences with internal and 5`- and 3`- end DNA modifications

| Name | DNA sequence with internal and 5` / 3`-end modifications |
|---|---|
| Antibody B* (antibody sequence) conjugated to IgG goat anti rabbit antibodies | TA CGC CCT AAG AAT CCG AAC TTG CAT TAC AGT CCT CAT AAG T /3`AmC3/ |
| DNA B1-Alexa Fluor 532 (antisense sequence) | 5`Acr ACT TAT GAG GAC TGT AAT GCT /3`Alexa Fluor 532/ |
| DNA B2-Alexa Fluor 532 (antisense sequence) | 5`Acr GTT CGG ATT CTT AGG GCG TAT /3`Alexa Fluor 532/ |
| Antibody B Cy5 (antibody sequence) conjugated to IgG goat anti rabbit antibodies | /5`Acr/ TA CGC CCT AAG AAT CCG AAC TTG CAT TAC AGT CCT CAT AAG T /3`AmC3/ |
| DNA B1-Cy5** (antisense sequence) | 5` ACT TAT G/iCy5/A GGA CTG /iCy5/TAA TGC A 3` |
| DNA B2-Cy5** (antisense sequence) | 5` AGT TCG G/iCy5/A TTC TTA /iCy5/G GGC GTA 3` |
| Antibody C Cy5* (antibody sequence) conjugated to IgG goat anti mouse antibodies | 5` GAC CCT AAG CAT ACA TCG TC TT GAC TAC TGA TAA CTG GAT TG /3`AmC3/ |
| DNA C1-Cy5** (antisense sequence) | 5` CAA TCC A/iCy5/G TTA TCA /iCy5/GTAGT CA 3` |
| DNA C2-Cy5** (antisense sequence) | 5` AGA CGA T/iCy5/G TATGCT /iCy5/TA GGG TC 3` |
| Modifications: Acr=Acrydite, AmC3=amino C3. Fluorescent dye modifications: iCy5=internal Cy5 dye, TMR=Tetramethylrhodamine, Alexa Fluor 532 | |

*Antibody B, Cy5-DNA B and Cy5-DNA C DNA sequences are equivalent to Antibody B and Antibody C sequences from Chen *et al.*[5]

**To avoid fluorophore interactions, the internal Cy5 dyes are separated by 7 base pairs, which also ensures that the fluorophores point into opposite directions. DNA sequences were ordered from Integrated DNA Technologies. All DNA sequences were HPLC purified and lypophilized. Antisense DNA oligos were re-suspended in TE buffer (10 mM Tris, 0.1 mM EDTA, pH 8.0) in nuclease-free water and kept frozen as aliquots with a concentration of 25 ng/µl at -20°C until use. Antibody sequences were conjugated to IgG (Rabbit) or IgG (Mouse) antibodies as indicated in Supplementary Table 2 using Solulink Antibody-Oligonucleotide All-In-One Conjugation Kit (Cat. No. A-9202-001). DNA labelled antibodies were kept as aliquots at 4°C.

**Supplementary Table 3**. Summary of Immunostaining and sample treatment in corresponding figures

| Figure | Specimen | Primary antibody (pre-expansion) | Secondary antibody / DNA label (pre-expansion) | Primary antibody (post-re-embedding) | Secondary antibody / DNA label (post-re-embedding) | Expansion protocol |
|---|---|---|---|---|---|---|
| Fig. 1b | Cos-7 cells | α-tubulin (ab1825), β-tubulin (T8328). | IgG Al532 (A-11002), IgG Al532 (A-11009). | | | proExM |
| Fig. 1f | Cos-7 cells | α-tubulin (ab1825) | IgG Al532 (A-11002) | | | unexpanded |
| Fig. 2b | Cos-7 cells | α-tubulin (ab1825), β-tubulin (T8328). | IgG Al532 (A-11002) | | | ExM-GA + Re-embedding |
| Fig. 2f | Cos-7 cells | β-tubulin (T8328), | Antibody C Cy5 DNA C1-Cy5, DNA C1-Cy5. | | | unexpanded |
| Fig. 2j | Cos-7 cells | α-tubulin (ab1825) | Antibody B, DNA B1-Al532, DNA B2-Al532. | | | ExM (DNA Al532) |
| Fig. 2n | Cos-7 cells | α-tubulin (ab1825), β-tubulin (T8328). | Antibody B Cy5, Antibody C Cy5 | | DNA B1-Cy5, DNA B2-Cy5, DNA C1-Cy5, DNA C1-Cy5. | DNA-Cy5 + Re-embedding |
| Fig. 3 | Cos-7 cells | α-tubulin (ab1825), β-tubulin (T8328). | IgG Al532 (A-11002) | α-tubulin (ab1825) | IgG Al532 (A-11002) | proExM + Re-embedding |
| Fig. 4a-c | Isolated centrioles | | | α-tubulin (ab1825) | Alexa Fluor 647 F(ab`)$_2$ (A-21246) | U-ExM + Re-embedding |
| Fig. 4e-g | Isolated centrioles | | | α-tubulin (ab1825) | HMSiR 647 F(ab`)$_2$ (A208-01) | U-ExM + Re-embedding |
| Fig 4h | Isolated centrioles | Poly-glutamylated tubulin (AG-20B-0020) | Alexa Fluor 647 F(ab`)$_2$ (A-21235) | | | unexpanded |

*MEA = cysteamine hydrochloride. Alexa Fluor dyes (Thermo Fisher) are abbreviated with Al. Antibodies are specified by the purchase order number of the respective supplier. A list of antibodies can be found under ‘Antibodies and labelling reagents` in the Online Methods section.

**Supplementary Table 4**. Summary of Immunostaining and sample treatment in corresponding supplementary figures

| Supp. Figure | Specimen | Primary antibody (pre-expansion) | Secondary antibody / DNA label (pre-expansion) | Primary antibody (Post-labeling) | Secondary antibody / DNA label (Post-labeling) | Expansion protocol |
|---|---|---|---|---|---|---|
| Supp. Fig. 2 | Cos-7 cells | $\alpha$-tubulin (ab1825), $\beta$-tubulin (T8328) | IgG Al532 (A-11002), IgG Al532 (A-11009). | $\alpha$-tubulin (ab1825) post-expansion | IgG Al532 (A-11002) post-expansion | proExM |
| Supp. Fig. 3 | Cos-7 cells | $\alpha$-tubulin (ab1825) | IgG Al532 (A-11002) | $\alpha$-tubulin (ab1825) post-expansion | IgG Al532 (A-11002) post-expansion | proExM + Re-embedding |
| Supp. Fig 4a | Cos-7 cells | $\alpha$-tubulin (ab1825) | IgG Al532 (A-11002) | | | unexpanded |
| Supp. Fig 4b | Cos-7 cells | $\alpha$-tubulin (ab1825) | Antibody B, DNA B1-Al532, DNA B2-Al532. | | | ExM (DNA Al532) |
| Supp. Fig. 6 | Cos-7 cells | $\alpha$-tubulin (ab1825) | Antibody B, DNA B2-Al532, DNA B2-Al532. | | | unexpanded |
| Fig. 9a-b | Isolated centrioles | | | $\alpha$-tubulin (ab1825) | Alexa Fluor 647 F(ab`)$_2$ (A-21246) | U-ExM + Re-embedding |
| Fig. 9c-d | Isolated centrioles | | | $\alpha$-tubulin (ab1825) | HMSiR 647 F(ab`)$_2$ (A208-01) | U-ExM + Re-embedding |

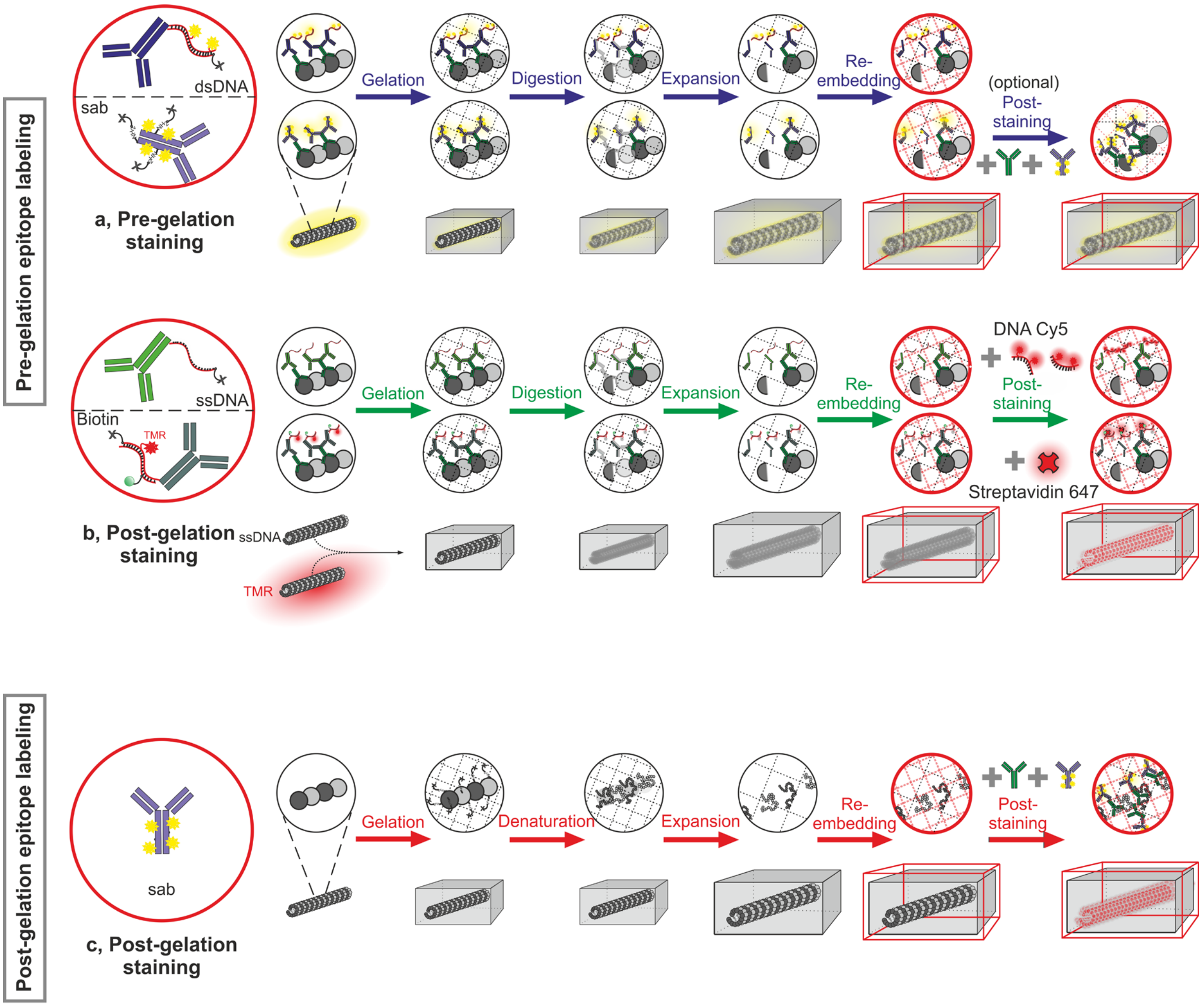


**Supplementary Figure 1**. **Schematic overview of pre- and post-labeling protocols used for Ex-SMLM**. The schematic is divided into two Ex-SMLM workflow strategies that mainly differ at the time of epitope labeling. The first strategy is to label epitopes before gelation, which in turn can be subdivided into protocols that introduce fluorophores before or after hydrogel formation. **a**,**b Pre-gelation epitope labeling (pre-labeling ExM) methods**. **a**, Pre-gelation epitope labeling (pre-labeling ExM). This workflow describes the use of DNA labeled secondary antibodies (dsDNA) used in the original ExM protocol as well as conventional fluorescent-dye conjugated secondary antibodies (sab) that are cross-linked into the polymer hydrogel network either by an acrydite DNA modification or through the linking of amine groups of the antibodies. After digestion with Proteinase K and expansion in water samples were re-embedded in an uncharged polyacrylamide gel, placed on a Bind-Silane treated coverslip, and optionally post-labeled (post-re-embedding) with the same primary and secondary antibodies for fluorescent signal amplification. **b,** Post-gelation labeling belongs also to the pre-labeling ExM methods because the linkage error is mainly determined by pre-labeling with antibodies. Ex-SMLM can also be performed with DNA modified secondary antibodies that are used for pre-gelation epitope labeling but fluorophores are introduced after gelation, digestion, expansion and re-embedding of the samples. Here secondary antibodies are modified with a single stranded DNA (ssDNA) that is directly incorporated into the hydrogel or a functionalized dsDNA carrying a Biotin insertion (for post-re-embedding staining with fluorophore conjugated biotin-binding streptavidin) and optional a TMR modification (for pre-expansion control imaging). **c**, **Post-gelation epitope labeling (post-labeling ExM).** Ex-SMLM can also be performed using ExM protocols like U-ExM where proteins themselves are anchored into the swellable gel matrix and immunostained post-expansion or post re-embedding using conventional primary and secondary antibodies. SMLM can be performed after expansion in pure water using the spontaneously blinking dye like HMSiR 647. Labeling after re-embedding enables also the use of *d*STORM dyes like Alexa Fluor 647 in photoswitching buffer.

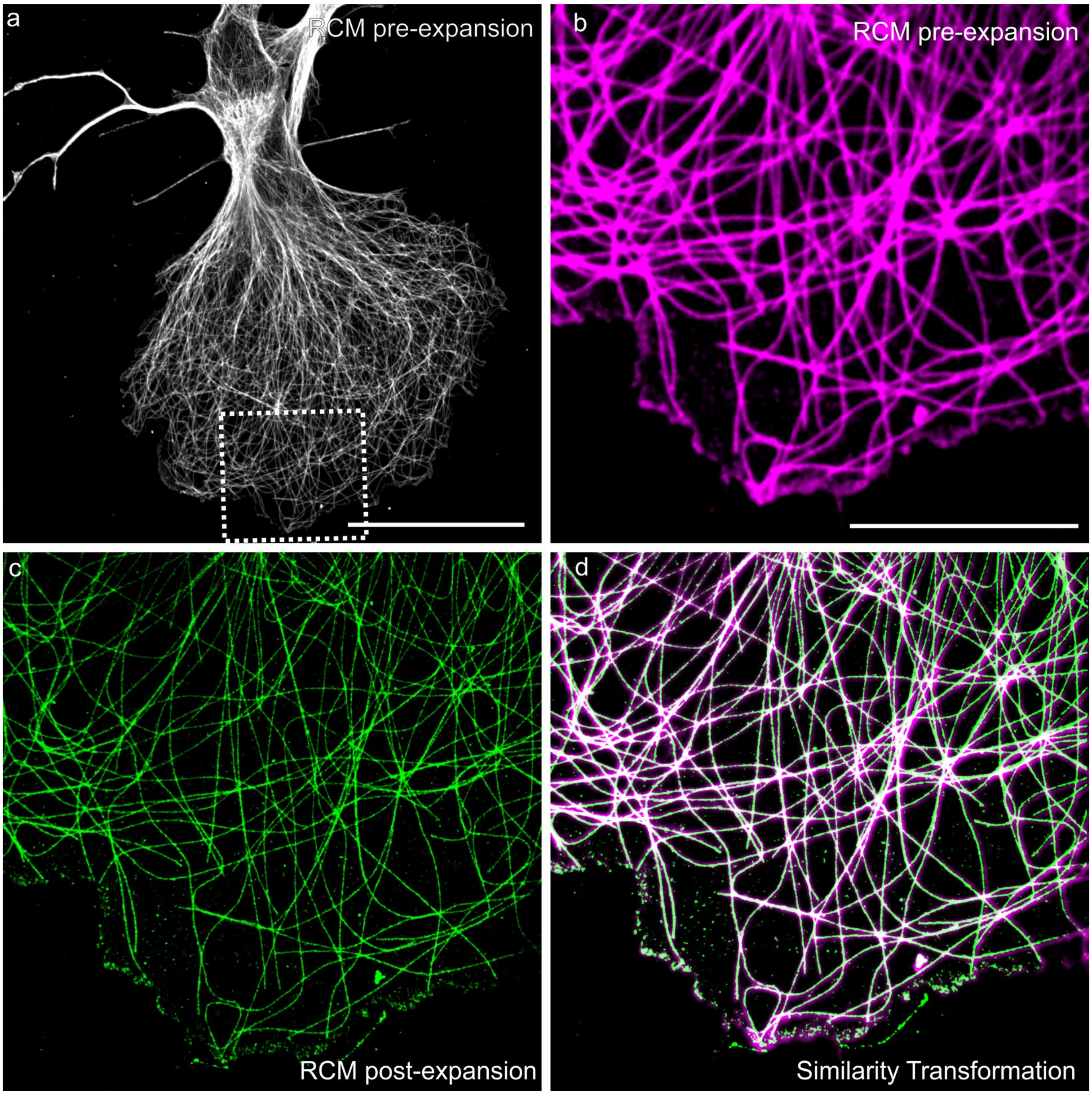


**Supplementary Figure 2. Expansion factor determination of pre- and post-expansion images. a,** Re-scan confocal microscopy (RCM)[2] image of unexpanded Cos-7 cells stained for tubulin with Alexa Fluor 532 conjugated secondary antibodies. **b,** Zoom-in on the highlighted region in (a). **c,** Post-expansion RCM image of the corresponding region shown in (b). **d,** Overlay of similarity transformed pre- and post-expansion RCM images shown in (b-c). An expansion factor of 3.84x was determined from the transformation parameters. Scale bars, 50 µm (a), 15 µm (b).

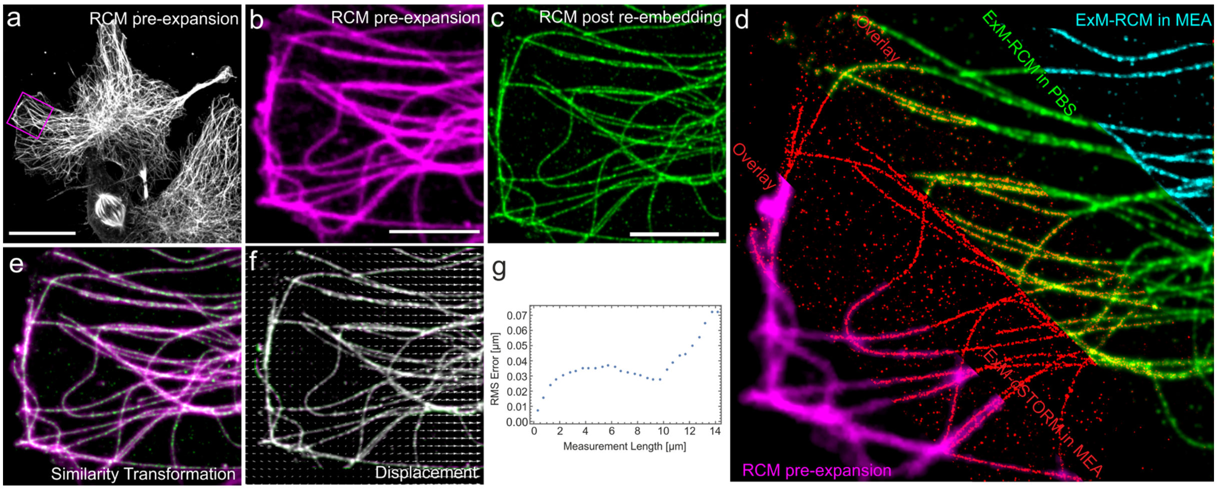


**Supplementary Figure 3. Expansion factor post re-embedding in different imaging buffer. a,** RCM image of α- and β-tubulin immunostained Cos-7 cells with Alexa Fluor 532 labeled IgG secondary antibodies. **b,** Zoom in on magenta boxed region in (a). **c,** RCM image of the expanded sample showing the corresponding area of (b) recorded in PBS (1x). The same monomer solution was used as in the experiments shown in Supplementary Fig. 2. **d,** Overlay of registered RCM and *d*STORM images taken either pre-expansion or post-re-embedding. Under each imaging condition the expansion factor after re-embedding determined by a similarity transformation resulted in an expansion factor of 3.1x registered to the pre-expansion RCM image. **e,** Overlay of pre-expansion RCM image (magenta) after alignment of the post re-embedding RCM image (green) acquired in PBS. **f,** Distortion vector field of data shown in (b) and (c). **g,** Root mean square error (RMSE) plot of data analyzed in (e). Scale bars, 25 µm (a), 5 µm (b), 15 µm (c).

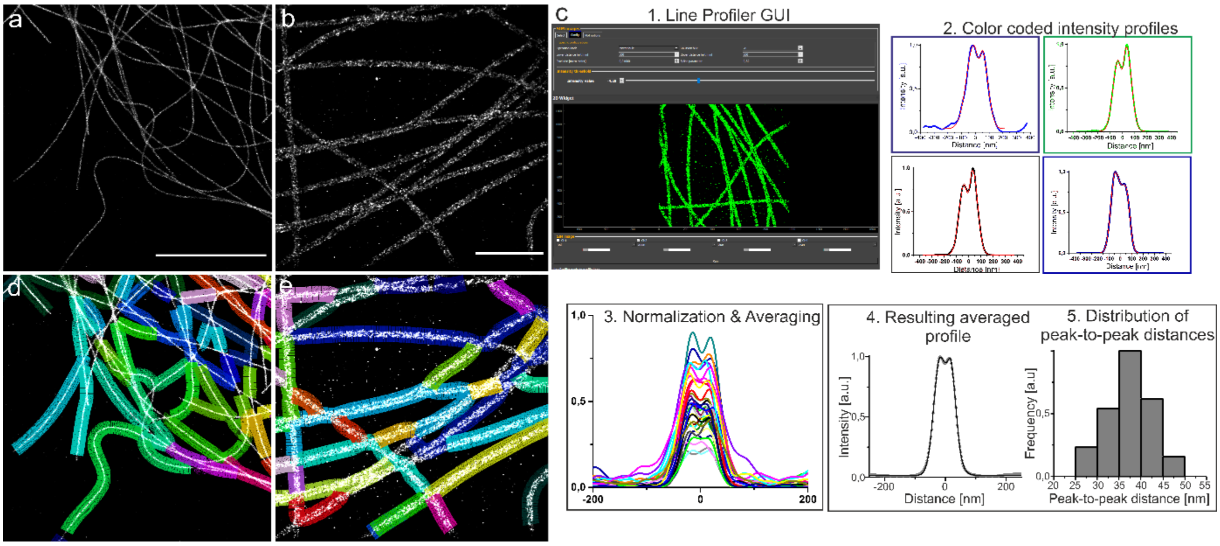


**Supplementary Figure 4. Analysis of microtubule cross-sectional profiles using LineProfiler. a-b,** Ex-SMLM input images of unexpanded Cos-7 cells labeled with antibodies against α-tubulin (a) and expanded Cos-7 cells labeled according to the ExM protocol using DNA trifunctional labels modified with Alexa Fluor 532 (ExM protocol) (b). **c, 1.** Graphical user interface of our automated image processing software Line Profiler (https://line-profiler.readthedocs. io/en/latest/)[3]. The software can be executed as .exe file and is able to load and visualize .tif images as shown in (a-b). Specific configurations as pixel size, Gaussian blurring and spline parameters can be adjusted for varying expansion factors, labeling densities and imaging modalities to optimize the automated detection of filaments. The functions Gaussian, Bi-gaussian, Tri-gaussian, cylinder projection and multi cylinder projection can be selected to fit the cross-sectional profiles along the filamentous structure. **2.** As an output the software gives color-coded cross-sectional profiles fitted to the selected functions and corresponding parameters as peak-to-peak distance or intensity values. An overview image with the analyzed segments **(d-e)** also shows the detected and analyzed filament segments. An examination of the analyzed segments is necessary to ensure cross-sectional profiles are set only at single filaments. Segments that exhibit crossing or several parallel running filaments must be excluded from the evaluation. **3-5.** Output txt. files of the manually chosen cross-sectional profiles can then further be analyzed in data analysis softwares like 'OriginPro' to get information about the averaged intensity profile (4.) and peak-to-peak distance distribution (5.).

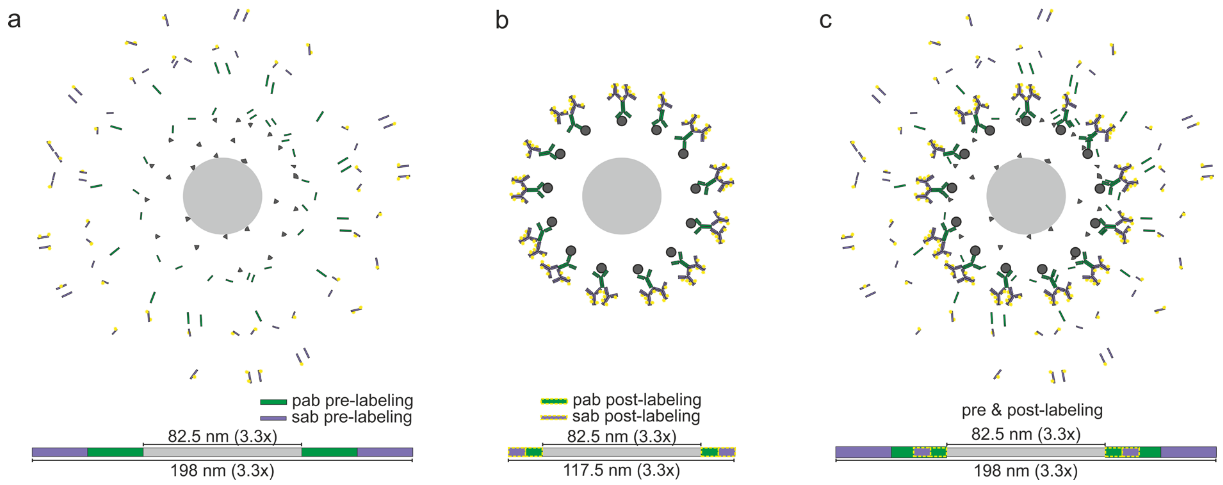


**Supplementary Figure 5. Schematic of inner and outer radius determination used for simulating cross-sectional profiles of expanded microtubules.** An exemplary schematic is shown that illustrates 3.3-fold expanded microtubules that are labeled pre-expansion with primary and secondary IgG antibodies as shown in Fig. 1a. Unexpanded microtubules decorated with primary and secondary IgG antibodies exhibit a diameter of 60 nm in electron microscopy experiments[1]. The total diameter after 3.3x expansion increases to 198 nm, which corresponds to a linkage error of ~58 nm. The inner diameter in which the labels (primary and secondary antibodies) are distributed after expansion is 82.5 nm determined by the 3.3x expansion of the 25 nm microtubule. The theoretically determined values (here shown for proExM and ExM-GA protocols) of the inner and outer diameter of the area in which the ExM labels are located after expansion were used to simulate cross-sectional profiles for different expansion methods and thus determine the molecular expansion factor (Supplementary Table 1 and Supplementary Note 1). **b,** Post-labeling ExM melts the linkage error down to 17.5 nm in a 3.3x expanded sample, which translates into a linkage error of ~ 5 nm for the unexpanded sample. Remarkably, the labeling efficiency increases substantially after expansion because of improved epitope accessibility (less steric hindrance of densely packed epitopes)[4]. Note that the antibodies are depicted to bind only to the outside of the expanded microtubule. An orientation of the antibodies towards the inside of the microtubule and binding of the antibodies in the inner area of the separated primary antibodies is also conceivable. **c,** Pre- and post-immunolabeling of microtubules with primary antibodies and fluorescently labeled secondary antibodies. Note that the pre-expansion signal is diluted ~ 36-fold ($3.3^3$) in three dimensions with an expansion factor of 3.3x. For the simulation of cross-sectional profiles (Fig. 2a and Supplementary Table 1) a two dimensional projection was used resulting in a ~11-fold ($3.3^2$) dilution of the signal after a 3.3x expansion.

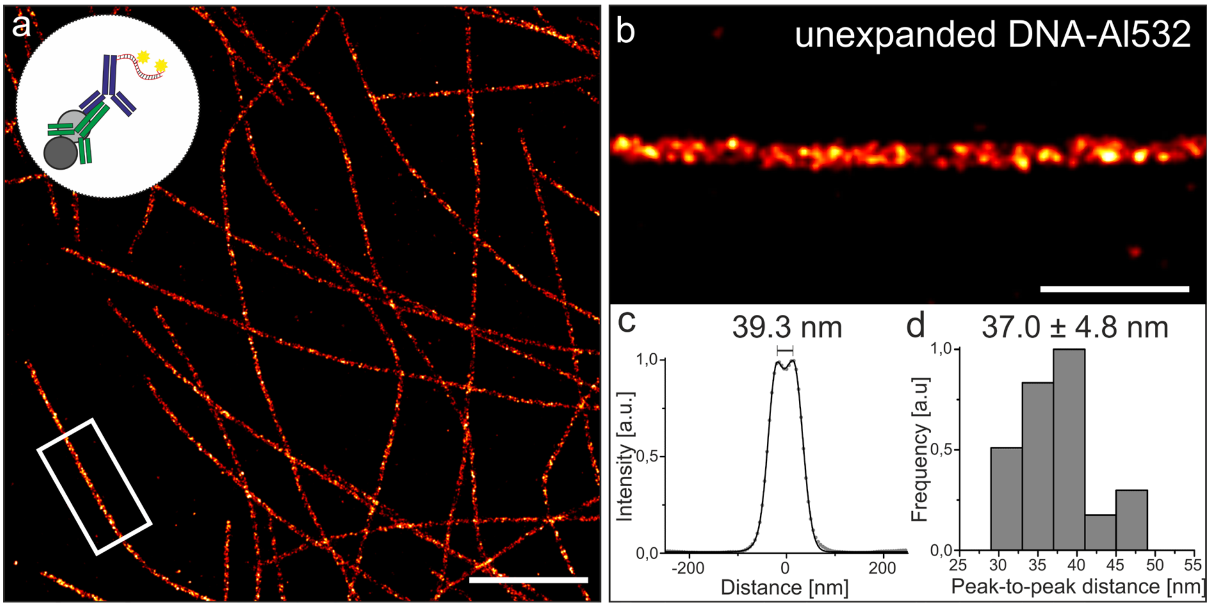


**Supplementary Figure 6. Cross-sectional profiles of unexpanded microtubules. a,** Unexpanded *d*STORM image of microtubule labeled with α-tubulin and ExM dsDNA label modified with Alexa Fluor 532 and anchored as dsDNA into the hydrogel polymer before expansion. **b,** Magnified view of highlighted region in (a). **c,** Averaged intensity profile of 34 microtubule segments with a total length of 98.1 µm (with individual lengths of 1.0-11.4 µm). **d,** Histogramed peak-to-peak distances of microtubule segments analyzed in (c) with a mean distance of 37.0 ± 4.8 nm (sd).Scale bars, 2 µm (a), 500nm (b).

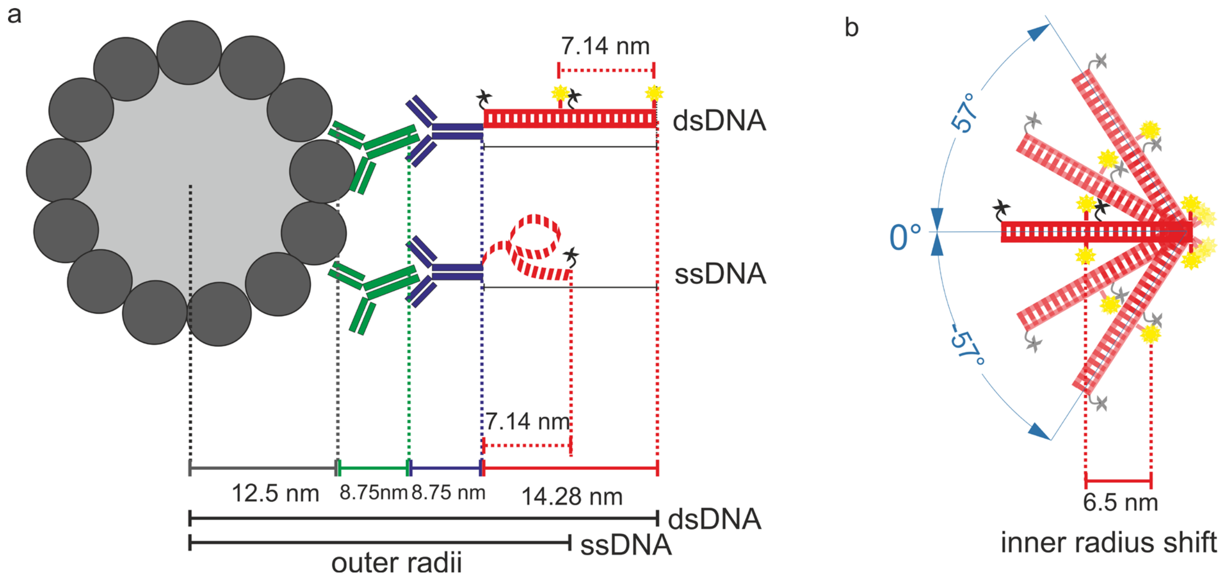


**Supplementary Figure 7. Schematic of DNA labels effecting the measured peak-to-peak distances. a,** Illustration of an immunolabeled microtubule filament consisting of 13 protofilaments (grey) labeled with primary antibody (green) and dsDNA or ssDNA modified secondary antibody. The scalebars show the broadening effect of the antibodies and DNA label on the measured intensity profile of the 25 nm thick microtubule. **b,** Spatial orientation of dsDNA label determined from experimental peak-to-peak distances of expanded microtubules using the ExM protocol with DNA trifunctional labels. The chart shows the distance shift of fluorophores that depends on the angle of the DNA. For the ExM protocol using DNA trifunctional labels the shift was determined to 6.5 nm.

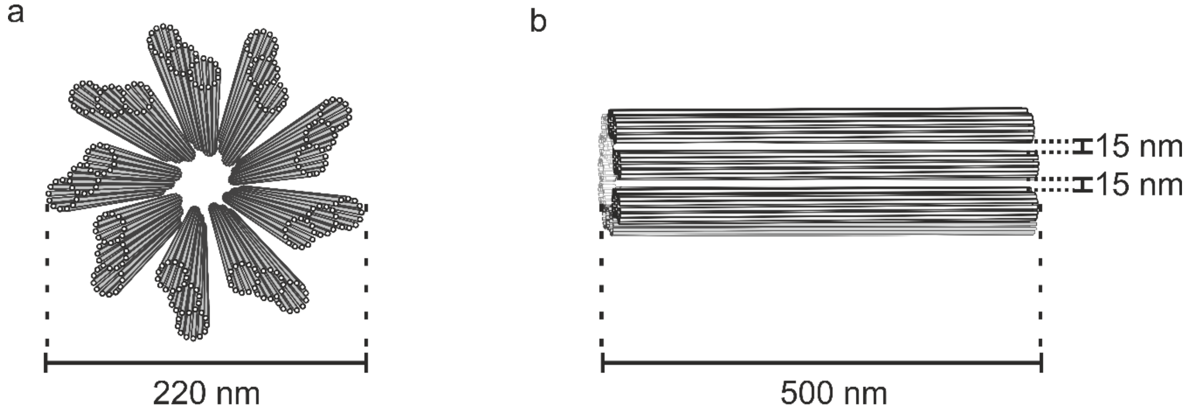


**Supplementary Figure 8. Model of centrioles. a,** Schematic of a centriole in frontal view showing the cylindrical arrangement of 9 microtubule triplets with a diameter of 220 nm. **b,** Schematic of a cerntriole in lateral view with a length of 500 nm and a distance of 15 nm between neighbouring microtubule triplets.

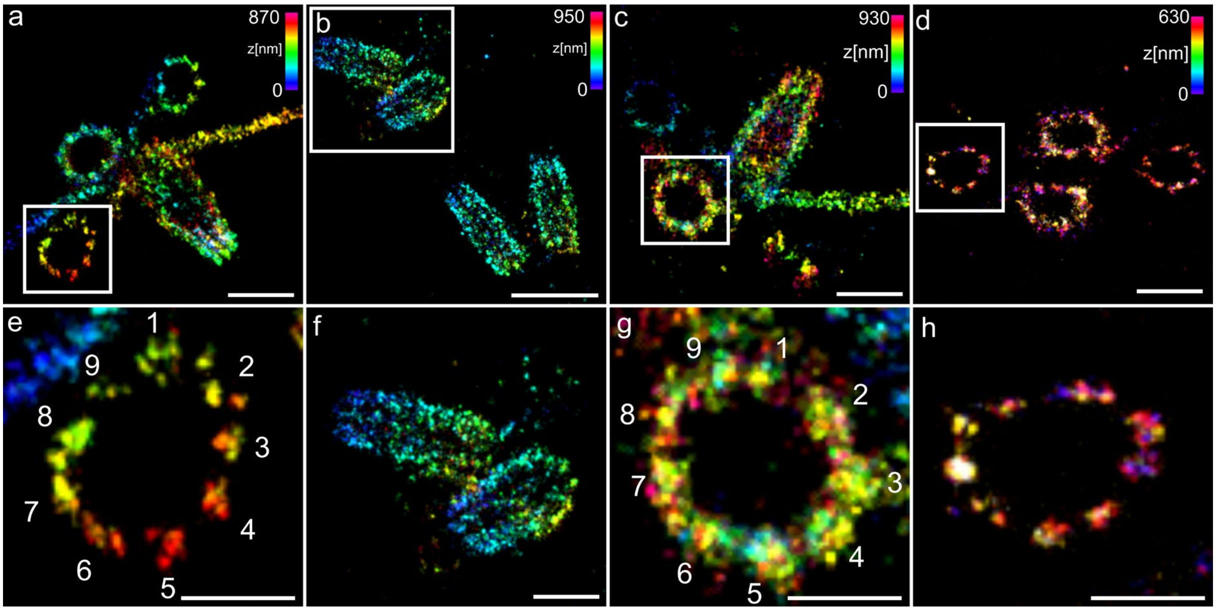


**Supplementary Figure 9. 3D Ex-SMLM of U-ExM expanded centrioles[4]. a,b,** 3D-*d*STORM images of *Chlamydomonas* centrioles immunostained with antibodies against α-tubulin and Alexa Fluor 647 post re-embedding imaged in photoswitching buffer. **c,d,** 3D-SMLM images of centrioles labeled with α-tubulin primary and HMSiR[5] conjugated secondary antibodies imaged in double-deionized water. **e-h,** Zoom-in on highlighted regions in (a-d) showing the 9-fold symmetry of centrioles at procentrioles (e, h) and a mature centriole (g) in frontal views and centrioles in lateral orientation (f). Scale bars, 1 µm (a), 2 µm (b), 1µm (c), 1µm (d), 500 nm (e), 1 µm (f), 500 nm (g-h).
